# Gene co-expression networks highlight key nodes associated with ammonium nitrate in sugarcane

**DOI:** 10.1101/2025.05.14.652917

**Authors:** Jorge Mario Muñoz-Pérez, Denis Bassi, Lucia Mattiello, Marcelo Menossi, Diego Mauricio Riaño-Pachón

## Abstract

Gene co-expression network analysis offers valuable insights into complex biological processes in plants. In this study, we investigated the molecular mechanisms underlying nitrogen utilization efficiency (NUE) in sugarcane by analyzing transcriptome data from two contrasting genotypes (responsive and non-responsive) across different leaf segments under varying nitrogen conditions. RNA-seq analysis of 48 samples identified 2,723 differentially expressed genes. The responsive genotype showed enrichment in carbon metabolism and defense response pathways under high nitrogen conditions, while the non-responsive genotype prioritized photosynthesis and stress response mechanisms. Differentially expressed genes were used to construct a co-expression network which revealed 199 distinct modules. 44 modules correlating to nitrogen availability and 20 to genotype. Notably, Module 20, containing a high proportion of MYB/MYB-related transcription factors, emerged as a potential key regulator of nitrogen response in both genotypes. Several modules showed correlation with important metabolic markers such as RUBISCO, chlorophyll, and PEPCASE. Additionally, we identified novel candidate genes, including uncharacterized transcripts with strong genotype-specific expression patterns. These findings provide potential targets for understanding and improving nitrogen use efficiency in sugarcane breeding programs.

## Introduction

Sugarcane is one of the most efficient crops in converting solar energy into sugars, capable of accumulating up to 0.7 M sucrose in mature stalks (Moore 1995). Despite this impressive efficiency, the theoretical maximum could be 70-80% higher under optimal conditions (Waclawovsky et al. 2010). Achieving this potential requires a deeper understanding of sugarcane physiology, particularly its photosynthetic performance. Unlike wheat and rice, where high photosynthetic rates have been a breeding focus (Makino 2011), sugarcane breeding has traditionally prioritized traits such as nutritional demands and stress resistance, even though enhanced photosynthesis could significantly boost productivity and biomass (Long et al. 2006; Marchiori et al. 2014).

Sugarcane has a C4 metabolism, which is highly regulated by the source/sink balance (Wardlaw 1990; McCormick et al. 2008). This metabolic pathway, coupled with environmental factors like CO_2_ concentration, light, temperature, and water availability, plays a crucial role in the plant’s photosynthetic activity. (Meinzer and Zhu 1998; Koonjah et al. 2006; Vu et al. 2006; De Souza et al. 2008; Yamori et al. 2014)

Among the nutrients, nitrogen (N) is particularly influential in photosynthesis and leaf development. It promotes protein synthesis, cellular growth, and the expansion of photosynthetic area, which in turn affects the production and translocation of photo-assimilates (Ryle et al. 1979; Theobald et al. 1998; Lawlor 2002; Vos et al. 2005; Kraiser et al. 2011; Leghari et al. 2016). In sugarcane, photosynthesis is closely linked to leaf N content, with a significant portion of N being mobilized from leaves to culms during the vegetative phase, negatively impacting the photosynthesis rate (Allison et al. 1997; Meinzer and Zhu 1998; Park et al. 2005). To enhance photosynthesis in sugarcane, new cultivars should be selected based on their ability to maintain high N levels in the leaves or to exhibit efficient photosynthesis nitrogen use efficiency (PNUE) (Sage et al. 2014).

Grass leaves provide an excellent model for studying C4 metabolism due to the developmental gradient along the leaf blade, where immature cells at the base transition to mature cells at the tip (Majeran et al. 2010; Sharpe et al. 2011; Mattiello et al. 2015; Bassi et al. 2018) However, the molecular mechanisms linking leaf development, nitrogen, and photosynthesis in sugarcane remain underexplored (Bassi et al. 2018).

Transcriptome analysis using high-throughput sequencing technologies is a powerful tool for investigating gene expression, structure, and regulation in crops and has enabled the rapid advancement in our understanding of various physiological and molecular processes in sugarcane (Garg and Jain 2013; Dong and Chen 2013; Dong et al. 2017). For example, transcriptomics has shed light on the mechanisms underlying sucrose accumulation (Huang et al. 2016; Hoang et al. 2017), lignin production (Vicentini et al. 2015), and the plant’s responses to both biotic (Brigida et al. 2016) and abiotic stresses (Gentile et al. 2013; Li et al. 2016; Diniz et al. 2020). Gene expression profiles at various developmental stages along leaf (Mattiello et al. 2015), the regulation of carbon partitioning (Correr et al. 2020), alternative splicing events related to the circadian clock (Dantas et al. 2019), in response to the hormone ethylene (Cunha et al. 2017), the pathways involved in cell wall biosynthesis (Hosaka et al. 2021) and association of gene expression with biomass content and composition (Hoang et al. 2018)

Gene co-expression network analysis has emerged as a valuable approach in crop research, offering a systems-level perspective on gene interactions (Rao and Dixon 2019). In sugarcane, this technique has been used to study Smut Resistance (Wu et al. 2022), transcription factors associated with cell wall biosynthesis (Ferreira et al. 2016), drought response (Li et al. 2022) and (Diniz et al. 2020), sugar and fiber accumulation (Perlo et al. 2022), response to mepiquat chloride in sugarcane (Chen et al. 2021), internode elongation (Chen et al. 2020), miRNAs and their role in nitrogen assimilation (Gao et al. 2022), among other studies. In this study, we applied gene co-expression network analysis to investigate the molecular mechanisms underlying differential nitrogen utilization efficiency (NUE) along the sugarcane leaf. This work represents the first comprehensive analysis of gene expression using co-expression networks in response to varying nitrogen levels across different leaf segments. Using two genotypes with contrasting NUE and two nitrogen concentrations, we identified differentially expressed genes (DEGs) and inferred co-expression networks to uncover genes and modules associated with nitrogen utilization. We hypothesize that genotypes with high NUE will exhibit distinct gene expression patterns, reflected in the co-expression networks, compared to those with low NUE.

## Methodology

### Plant material

We used RNASeq data generated during the same experiment as (Bassi et al. 2018), but that has not been published until now. We describe briefly the experimental setup in the following but we refer the reader to (Bassi et al. 2018) for further details. Contrasting sugarcane genotypes, RB975375 (responsive, R) and RB937570 (nonresponsive, NR), were selected based on nitrogen-use efficiency (NUE) screening, involving a pool of 20 genotypes exposed to three nitrogen concentrations (10, 90, and 270 mg of N per kg of sand) using ammonium nitrate as nitrogen source as described by (Robinson et al. 2008). The culms were sprouted in greenhouse trays filled with vermiculite. After three weeks, the plantlets were transferred to plastic pots containing 3.4 kg of washed sand and maintained in a greenhouse at a constant temperature of approximately 28°C. The nitrogen was applied at two concentrations: low N (10 mg N/kg sand) and high N (270 mg N/kg sand). Applications occurred three times at 15-day intervals. The experimental design was completely randomized, with three biological replicates per genotype and treatment combination. Each biological replicate consisted of a single plant, and samples were collected from four distinct leaf segments per plant. This resulted in a total of 48 samples. Leaf segments were harvested from leaf +1 (the first fully expanded, photosynthetically active leaf) of three-month-old plants, one month after the final N application. The leaf blade was divided into four developmental zones: Base Zero (B0): The first 2 cm of the leaf base, representing immature tissue. Base (B): Middle region of the first third, transitioning from cell division to elongation. Middle (M): Middle region of the second third, with mature photosynthetic cells. Tip (P): Final third of the leaf. Segments were collected between 10 AM and 2 PM to minimize diurnal variation, immediately frozen in liquid nitrogen, and stored for downstream analyses.

### RNA extraction, library construction, and sequencing

Total RNA was extracted from three independent replicates for each sample and treatment using Trizol (Invitrogen, USA), with an additional sodium acetate/ethanol precipitation step to enhance purity. RNA quality and concentration were assessed using gel electrophoresis, a NanoDrop spectrophotometer (Thermo Fisher Scientific, USA), and a Bioanalyzer (Agilent Technologies, USA). Only RNA samples with a minimum RNA Integrity Number (RIN) of 7 were used for library construction. A total of 48 libraries were prepared using the TruSeq Stranded mRNA Sample Prep Kit (Illumina), which enriches poly-A-containing transcripts and retains strand information. Clusters were generated on a c-Bot (Illumina), and paired-end sequencing was performed on a Hi-Seq 2500 platform (Illumina) with the TruSeq SBS Kit v3 – HS (Illumina). Sequencing was conducted at the LACTAD Facility (University of Campinas, Campinas, Brazil). Raw reads can be found under NCBI bioproject PRJNA1176579.

### RNASeq data pre-processing, assembly

Short-reads were pre-processed using FastQC (Andrews 2010), multiQC (Ewels et al. 2016), BBDuk2 v35.85 (Bushnell 2014) and Trimmomatic v0.38 (Bolger et al. 2014) to remove low quality regions, remaining adaptor sequences, chloroplast and mitochondria reads, rRNAs, and rat/mouse contamination. All the reads from the same genotype were used to generate a de novo transcript assembly with Trinity 2.14.0 (Grabherr et al. 2011) in strand-specific mode and setting the kmer value to 25 bp and minimum transcript length of 201. The resulting genotype-specific transcript assemblies were joined into a comprehensive transcriptome.

### Transcript quantification and grouping

Transcript abundance was quantified with Salmon v1.10.2 (Patro et al. 2017) using the assembled transcriptome as reference. Due to the high ploidy and alternative splicing isoforms present in sugarcane, some of the assembled transcripts could be quite similar and thus share many sequencing fragments, so we decided to cluster them based on the equivalence classes inferred by Salmon v1.10.2 (Patro et al. 2017) using Terminus (Sarkar et al. 2020). The quantification at the transcript level generated by Salmon was summarized at the level of groups inferred by Terminus, and we will refer to these groups as “transcript groups” hereafter.

### Functional annotation of the *de novo* transcriptome

Predicted peptides using TransDecoder v5.5.0 (Haas, BJ. https://github.com/TransDecoder/TransDecoder) were annotated with the Trinotate pipeline v3.3.2 (Bryant et al. 2017), which includes sequence similarity searches against the Swissprot v2020_06 database with BLASTX and BLASTP v2.8.1 (Altschul et al. 1990), prediction of signal peptides using SignalP v5.0b (Teufel et al. 2022), prediction of transmembrane regions using TMHMM v2.0c (Krogh et al. 2001), identification of ribosomal genes with RNAmmer v1.2 (Lagesen et al. 2007), Gene ontology (GO) (Ashburner et al. 2000; Gene Ontology Consortium et al. 2023), Transcription associated proteins (TAPs) domains were identified using Hmmer v3.3.2 (Eddy 2011) and Pfam v34 (El-Gebali et al. 2019). Protein domains were classified into TAPs families using PlnTFDB (Riaño-Pachón et al. 2007).

### Quantification, quality control and differential gene expression analyses

Quantification by Salmon v1.10.2 (Patro et al. 2017) was imported into R using the tximport package (Soneson et al. 2016). We transformed the raw counts using the *vst* function from DESeq2 (Love et al. 2014) which allows us to stabilize the variance across the range of the mean, handle zero counts (Anders and Huber 2010) and provides a more suitable basis for heatmap representation and co-expression analyses. We conduct PCA analysis using the *ploPCA* function from DESeq2 which uses the top 500 transcript groups with highest row variance to assess the grouping of the samples according to the experimental design. To evaluate the effect of Nitrogen levels on gene expression we compared the expression profiles in the B0 (Base zero), B (base), M (Middle), and P (Tip) regions of the leaf from sugarcane plants grown under 10 and 270 mg of N per kg of sand. Using the raw counts matrix with normalizing factors computed by DESeq2, we identified differentially expressed transcript groups in eight nitrogen availability contrasting conditions (Table 1). We used DESeq2 (Love et al. 2014) with params *altHyphothesis = graterAbs* and *lfcThreshold = 1* in order to get differentially expressed transcript groups two times above/under the background expression level.

**Table 1.** Contrasts for differential expression analysis. Nitrogen responsiveness = NR: non-responsive genotype (RB937570), R: responsive genotype (RB975375), Leaf segments = B0: Base 0, B: Base, M: Medium, P: tip, Nitrogen level = 10: 10 mg of N per kg of sand, 270: 270 mg of N per kg of sand. We carried out eight statistical contrasts, in each of them genotype and leaf segment remained constant and only Nitrogen level varied.

| Contrast |
| --- |
| NR-B0-10 vs NR-B0-270 |
| NR-B-10 vs NR-B-270 |
| NR-M-10 vs NR-M-270 |
| NR-P-10 vs NR-P-270 |
| R-B0-10 vs R-B0-270 |
| R-B-10 vs R-B-270 |
| R-M-10 vs R-M-270 |
| R-P-10 vs R-P-270 |

### Inference of co-expression network and modules

All transcript groups detected as differentially expressed in at least one of the contrasts were kept for network inference, and counts after the variance stabilizing transformation were used as expression values for network inference, analysis and heatmap visualization. We filtered out transcript groups with more than a half missing entries and zero variance with the function *goodSamplesGenes* from WGCNA (Langfelder and Horvath 2008). Pearson correlation coefficient between each pair of genes was computed with the *fastCor* function from HiClimR R package (Badr et al. 2015). Only gene pairs with |cor| > 0.90 were kept in the network. Clusters, also called modules of co-expressed transcript groups, were identified with the Markov cluster algorithm MCL (Van Dongen 2000) with an inflation value of 1.8. We estimated the Kullback-Leibler divergence between our graph an several random graph models: Erdős-Rényi (Erdos and Rényi 1960), Small-world (Watts and Strogatz 1998) and scale-free (Barabási and Albert 1999) as implemented in StatGraph R package (Takahashi et al. 2012)

### Functional attribution to gene sets and correlation analysis

In order to identify the most relevant biological functions in gene sets we carried out overrepresentation tests (Fisher’s exact test adjusted p-value < 0.05) of Biological Processes from the Gene Ontology using the TopGO R package (Alexa and Rahnenführer 2009). The gene sets used comprised the differentially expressed transcript groups in each of the contrasts and the co-expression modules. p-values were adjusted with the Bonferroni correction for multiple testing. We used the eigengene of each module which tends to get only one representative expression pattern of a set of transcript groups using the first principal component of their expression matrix (Zhang and Horvath 2005). Using this representative expression pattern for each module called eigengene we were able to calculate the correlation of modules with genotype, nitrogen availability and several metabolites determined previously for the exact same samples (Bassi et al. 2018) using the Spearman correlation coefficient.

## Data availability

Data is available via Figshare: https://figshare.com/projects/NitrogeGene_co-expression_networks_highlight_key_nodes_associated_with_ammonium_nitrate_response_in_sugarcane/242225, including transcript group expression values per sample, and transcript assemblies RNASeq datasets are available under the NCBI’s BioProject number PRJNA1176579

## Results and discussion

### Transcript assembly, annotation, quantification and exploratory analyses

We assembled 3,835,051 transcripts. Processing with Terminus to exploit equivalence classes resulted in 2,480,471 transcript groups, from which 1,116,882 (45%) encoded for at least one protein. 873,612 transcript groups were assigned to 24,063 GO terms, and the 2,502,895 deduced proteins were identified as belonging to 6,934 Pfam protein families.The PCA analysis indicates that differences in genotype explain 93% of the variance and only 2% of the variance is explained by different nitrogen levels and developmental stages, (Figure 1A). Within genotypes, the samples were grouped according to leaf segment and N condition. (Figures 1B and 1C)

**Figure 1.**
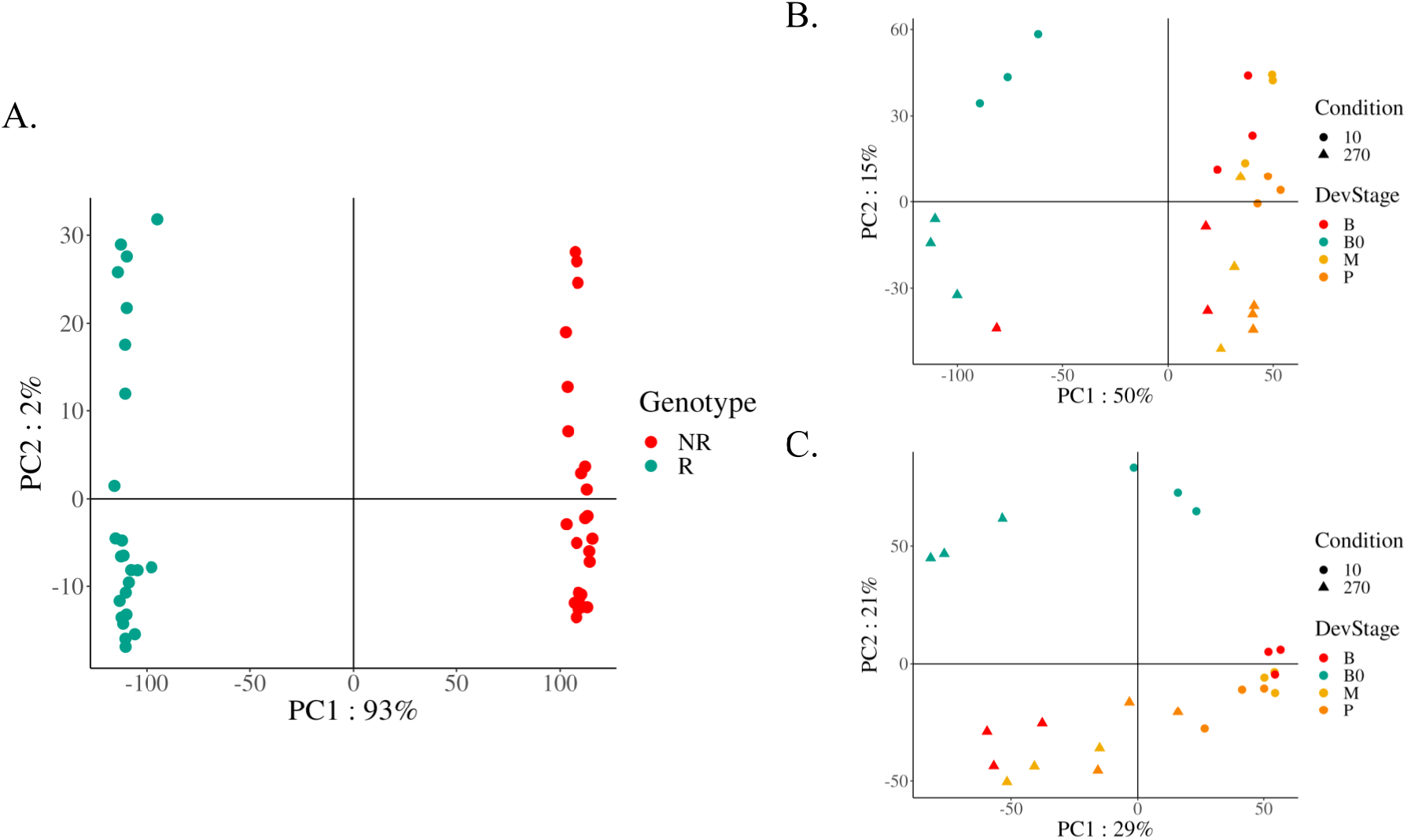
Principal component analysis representing the top 500 transcript groups by row variance in each sample. A). All genotypes. B). Samples from the Nitrogen responsive (R) genotype. C) Samples from the non responsive (NR) genotype

### Different responses to high nitrogen levels for sugarcane genotypes

From 2,480,471 transcript groups, we identified 2,723 differentially expressed (DEGs) (p < 0.01 and |log fold change| > 1) across at least one condition. The NR genotype showed a higher number of up-regulated transcript groups than the R genotype except for the tip of the leaf (P) in the R genotype which exhibited the greatest number of DEGs compared to all other conditions. Most DEGs were unique for each condition for both genotypes, suggesting different processes occurring in leaf segments. However, 21 transcript groups in the NR genotype were consistently up-regulated across all leaf segments in NR genotype (Figure 2). These transcript groups were enriched in response to ammonium ion GO:0060359 (p < 0.01).

**Figure 2.**
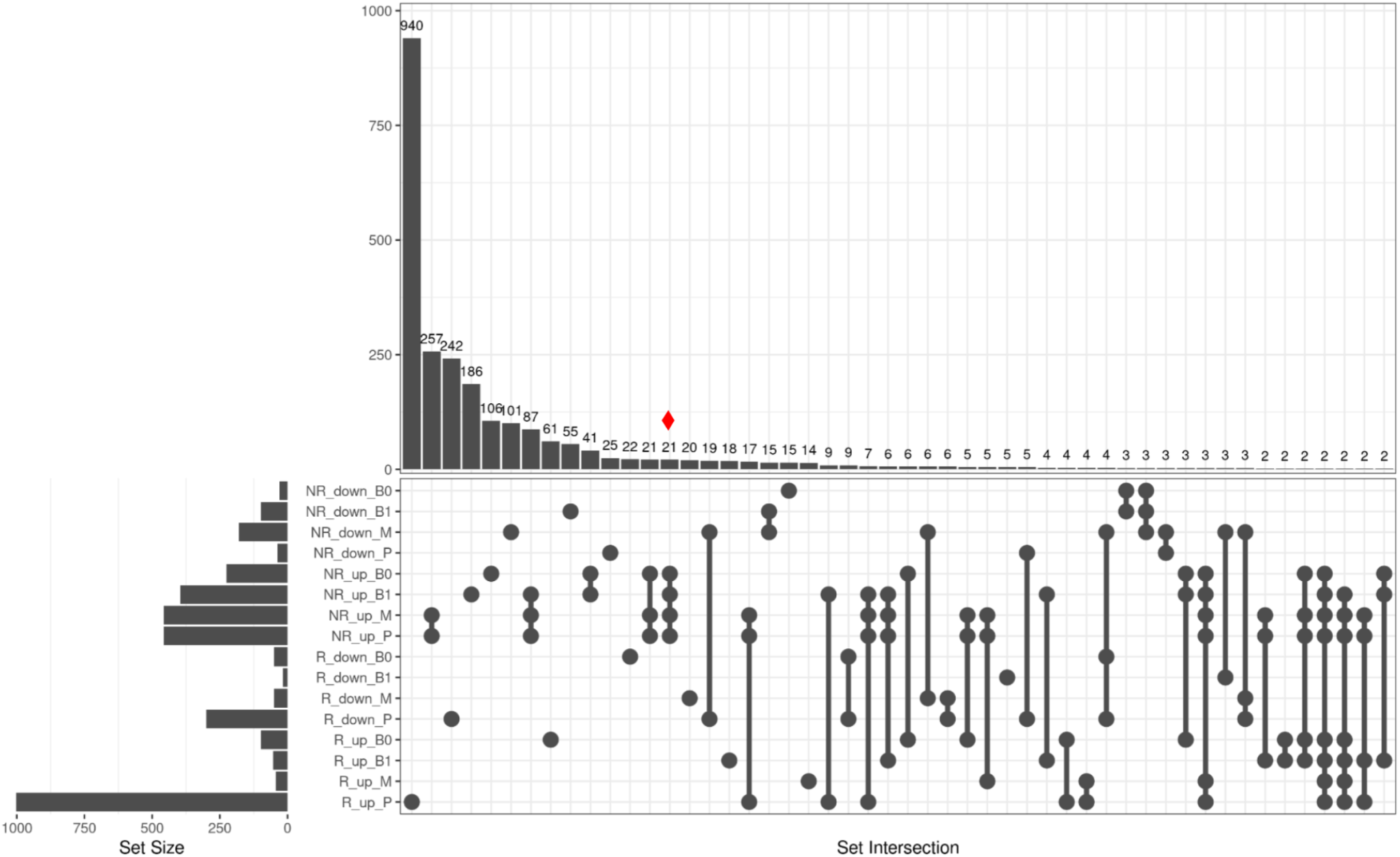
Sets of differentially expressed transcript groups; shared and unique across the different conditions. Up relates to the transcript groups overexpressed in high nitrogen availability. Down relates to transcript groups overexpressed in low nitrogen availability. Red diamond indicates a set of transcript groups always up-regulated in NR genotype in high nitrogen availability irrespective of the leaf segment.

In the responsive genotype, most regulated transcript groups respond to high nitrogen availability, specially in the tip of the leaf (P segment, R_up), the most relevant terms are related to chitin catabolic process, amino sugar catabolic process, and glucosamine-containing compound catabolic process (Figure 3A) This suggests a strong metabolic focus on breaking down complex sugars, likely reflecting a response to nitrogen availability by enhancing carbon metabolism. In contrast, the non-responsive genotype in high nitrogen availability (NR_up) shows photosynthesis (light reaction), cellular response to iron ion starvation and chitin catabolic process as the most relevant terms (Figure 3B). These processes indicate a focus on energy production and stress response to iron, highlighting a more resource-oriented reaction rather than a defensive one. Interestingly NR_up is enriched in photosynthesis terms, while metabolomics analyses showed more total chlorophyll in R_up along the entire leaf (Bassi et al. 2018). Additionally we found 47 chitinases in R_up and 36 in NR_up. They are frequently found in responses to biotic and abiotic stress (Vaghela et al. 2022) and N fertilization enhances chitinase levels (Dietrich et al. 2004). For the responsive genotype in low nitrogen availability (R_down), the most relevant terms are regulation of metal ion transport, cellular response to metal ions, and regulation of ion transport (Figure 3C). This suggests a heightened regulatory response to metal ion availability and homeostasis under low nitrogen, potentially as a compensatory mechanism for maintaining nutrient balance. The non-responsive genotype under low nitrogen availability (NR_down) similarly shows a strong focus on regulation of potassium ion transport and metal ion transport, indicating a shared importance of ion regulation under nitrogen stress. The shared terms across genotypes (ion transport and response to metal ions) point to a common stress response mechanism. While both genotypes respond to nitrogen conditions with shared processes, the responsive genotype (R) prioritizes metabolic and regulatory pathways, especially related breakdown of compounds with complex sugars and gene expression regulation. In contrast, the non-responsive genotype (NR) under high nitrogen emphasizes energy production and stress responses, while under low nitrogen it activates metabolic and catabolic pathways related to specialized compounds.

**Figure 3.**
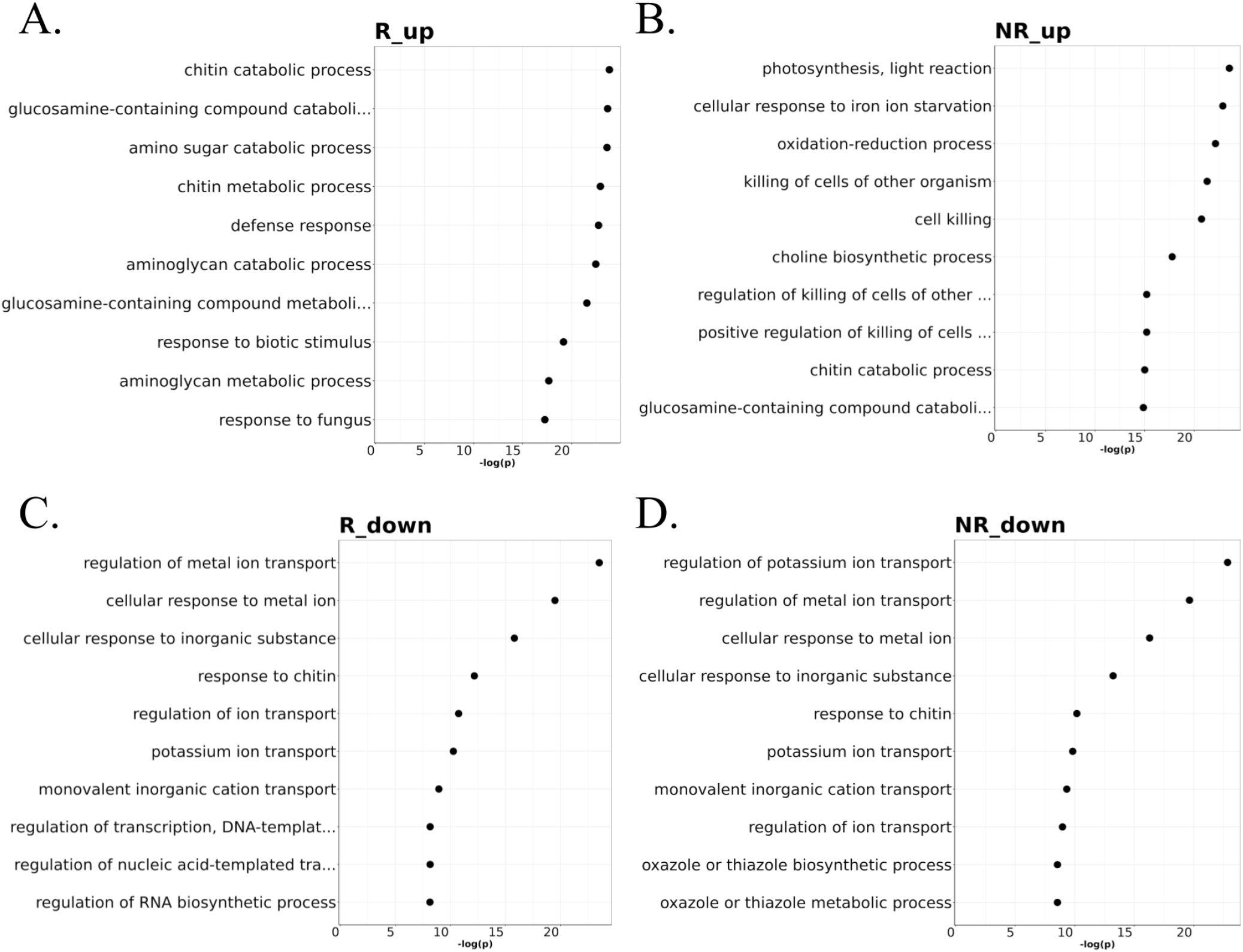
Top 10 functional enrichment of DEGs irrespective of leaf segment. A) Over expressed transcript groups in high nitrogen availability in the responsive genotype. B) Over expressed transcript groups in high nitrogen availability in the non-responsive genotype. C) Over expressed transcript groups in low nitrogen availability in responsive genotype. D) Over expressed transcript groups in low nitrogen availability in non-responsive genotype.

### Gene co-expression network inference and clustering

From the set of 2,723 differentially expressed transcript groups, we inferred a gene co-expression network (|cor| > 0.9) yielding 1,109 nodes, 3,699 edges and 110 connected components, the giant component comprising 56.4% of the nodes of the network (Figure 4). The Kullback-Leibler divergence between our graph and a Barabási-Albert with the same number of nodes and a parameter equal to 1.3 was only of 0.08 indicating that our graph is well explained by a Barabási-Albert model. This indicates a pronounced clustering tendency and suggests a non-random organization of co-expressed nodes within our network We retrieved 199 modules with at least 2 nodes on them, with the biggest module, module one, containing 73 nodes (6.58% of all nodes in the network), followed by module two with 44 nodes (3.97% of all nodes in the network). 28 modules (14.07% of all modules in the network) had more than 10 nodes. This diversity in module sizes may reflect distinct functional categories or regulatory pathways within the co-expression network.

**Figure 4.**
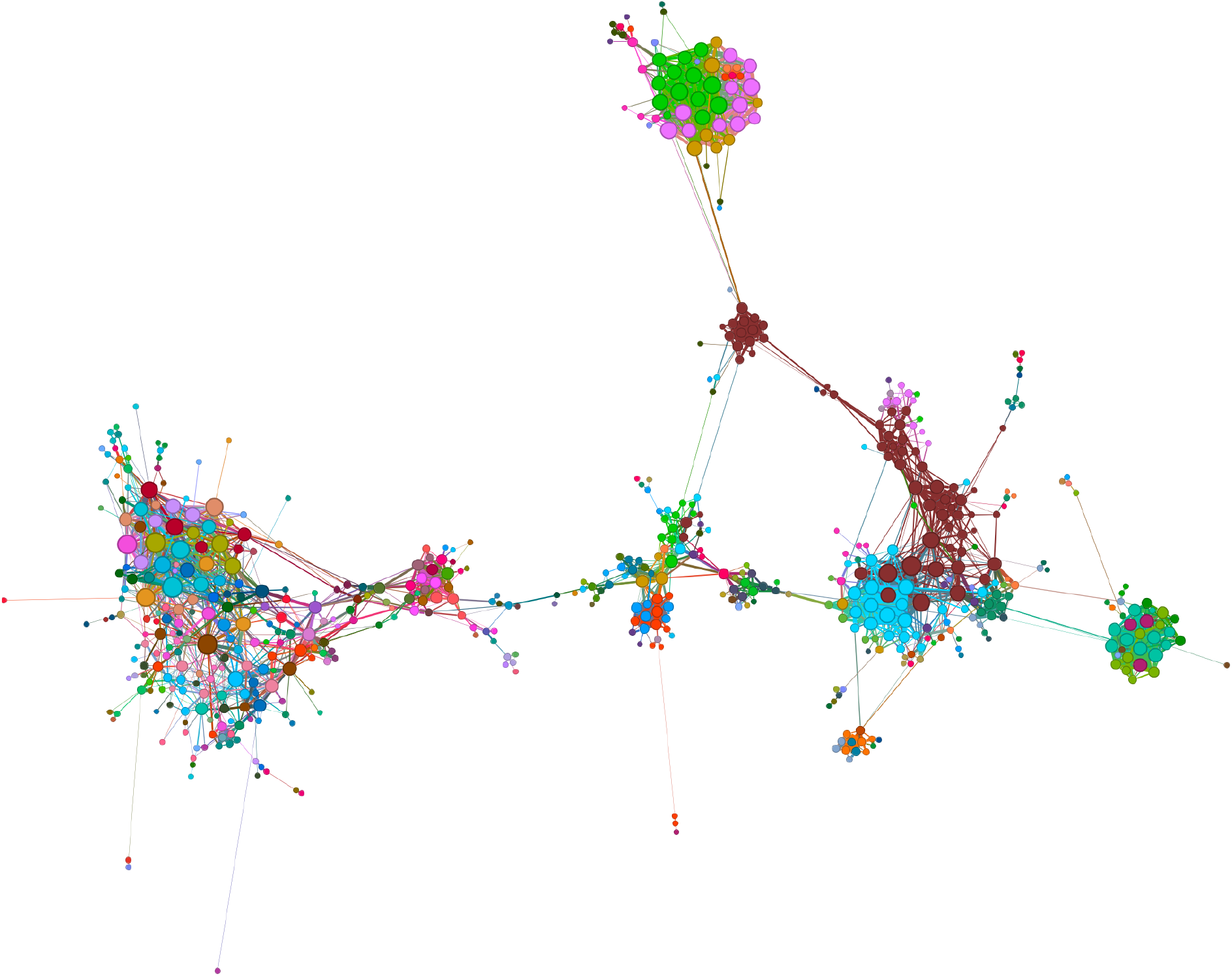
Network gene co-expression visualization. Same colors indicate the same modules. Node diameter is proportional to the degree.

For all modules, we got a list with the GO overrepresented terms for Biological process (BP), p-value < 0.01. Additionally, correlation analysis between modules, genotype and nitrogen availability was done to detect responsive modules for nitrogen levels and genotype

### Physiological/Biochemical traits associated with co-expression modules

We use the physiological/biochemical traits reported by Bassi et al. (2018) for the same samples, to look for correlation between these traits and the co-expression modules identified here. There were no modules correlated to genotype and nitrogen level at the same time. However there were 44 modules correlated to nitrogen level (|correlation| > 0.7, Bonferroni < 0.05) and 20 correlated to genotype (Figure 5). Among the modules correlated with the genotype we found several interesting modules, as described below:

**Figure 5.**
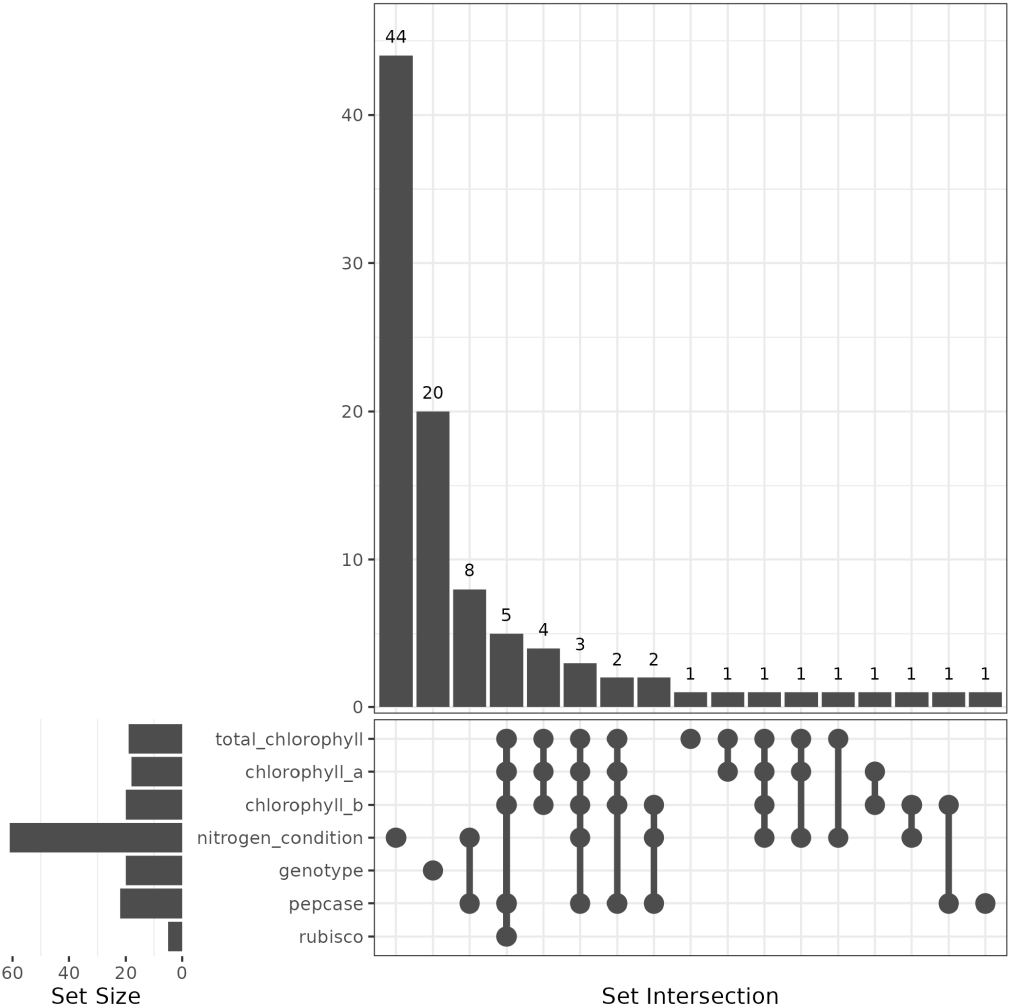
Sets of correlated modules with genotype, nitrogen condition and several metabolites.

#### Module 10

Focus on cellular energy metabolism, particularly emphasizing photosynthesis, glycolysis, and ATP production. Additionally, it is involved in cellular responses to cytokinins and the organization of organelles. The inclusion of enriched GO terms related to monosaccharide metabolism and oxidation-reduction processes reinforces its role on cellular energetics and metabolic regulation. The transcript groups in this module have a larger expression level in the NR genotype, within this genotype it is also seen a difference between the two nitrogen levels (Figure 6A)

**Figure 6.**
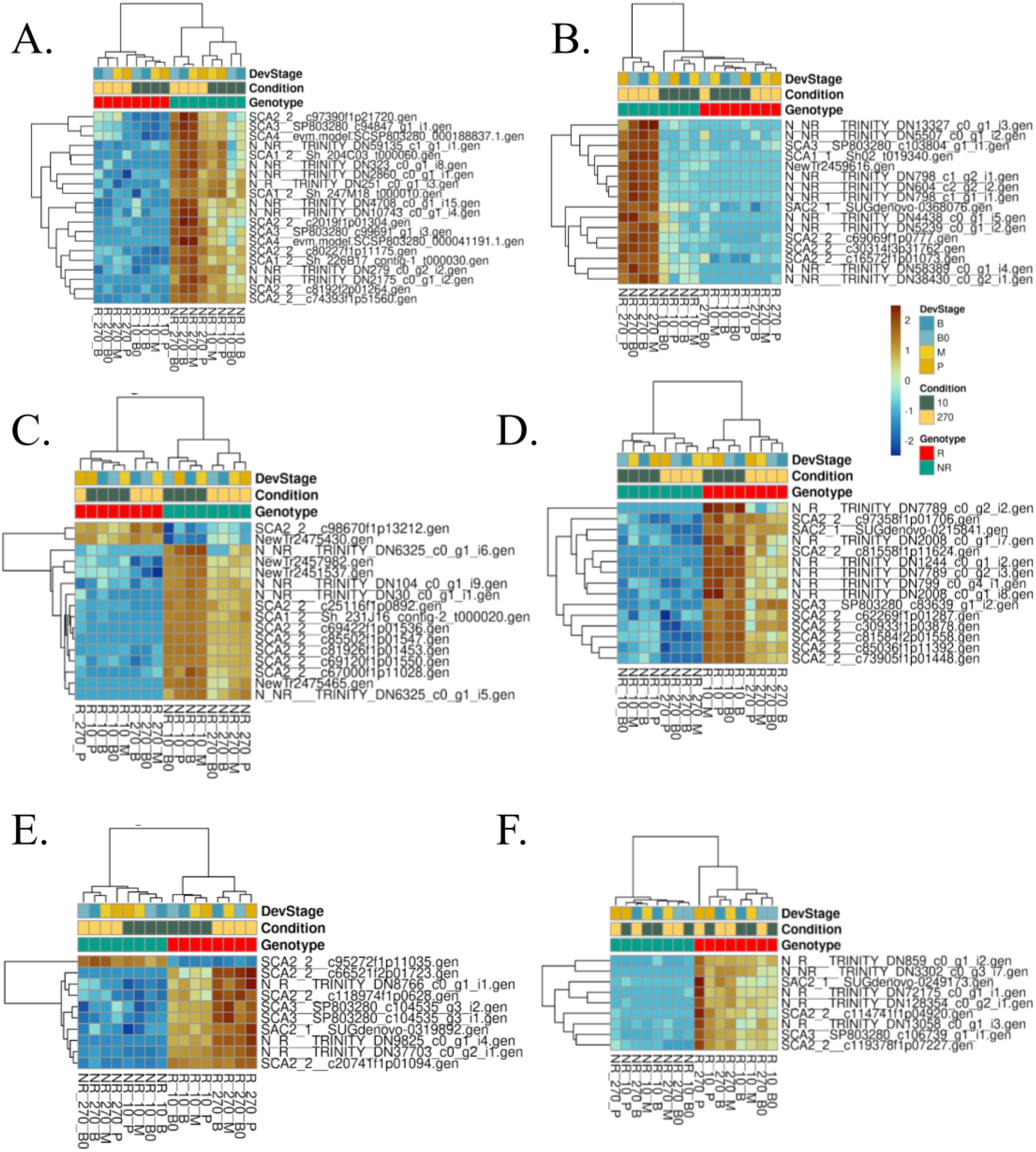
Z-score expression values of selected modules. A) Module 10. B) Module 13. C) Module 15. D) Module 17. E) Module 24. F) Module 29.

#### Module 13

Enriched in alpha-amino acid metabolic process, plant organ development, root system development, ammonium ion metabolic process, choline metabolic process this module was strongly activated only in the NR genotype at high nitrogen availability (Figure 6B), maybe pointing differences in the nitrogen usage between both genotypes at high nitrogen availability.

#### Module 15

Involved in essential cellular processes related to ion transport, gene expression regulation, biosynthesis of biomolecules, and cellular responses to various stimuli. The focus on ion transport and metabolic processes suggests roles in maintaining cellular homeostasis, suggesting a role in adapting to environmental changes, including nutrient availability and chemical signals. Remarcably this module had two genes activated only in R genotype: SCA2_2_c98670f1p13212.gen and NewTr2475430.gen (Figure 5C). The first gene is annotated as phosphoenol pyruvate carboxylase (PEPCASE), which might be one of the reasons that makes photosynthesis more efficient in the R genotype (Bassi et al. 2018). The second gene could not be annotated making it an attractive candidate for functional genomics analysis.

#### Module 17

Enriched GO terms suggests involvement in fundamental cellular processes related to responding to stimuli, regulating gene expression, synthesizing macromolecules, and orchestrating developmental processes. The presence of terms related to chromatin organization and protein modification suggests a role in epigenetic regulation, which can have deep effects on gene expression and cellular phenotypes. This module is strongly activated in the R genotype and less activated in the high-level nitrogen condition, compared to the low-level nitrogen condition (Figure 6D). The gene N_R__TRINITY_DN7789_c0_g2_i2.gen is grouped alone in the heatmap and has a response strongly dependent on the R genotype and the nitrogen level, being another gene with unknown function.

#### Module 24

Appears to be involved in cellular processes related to protein regulation, amino acid metabolism, and plant defense responses including non photochemical quenching and energy quenching, These processes are related to dissipating excess energy in photosynthetic organisms, especially under stress conditions. The presence of terms like protein modification, dephosphorylation, and macromolecule modification suggests a role in fine-tuning cellular activities and responses to environmental stimuli, including pathogen attacks and stress conditions. The focus on amino acid metabolism, particularly alanine and pyruvate family amino acids, may indicate metabolic adjustments in response to stress or nutrient availability. This module was heavily activated in the R genotype, particularly under the high-level nitrogen condition, except for the gene SCA2_2__c95272f1p11035.gen. This gene is annotated as Chlorophyll a/b binding protein CP26, which is deactivated in R genotype and activated in NR genotype (Figure 6E).

#### Module 29

This module is highly activated only in R genotype (Figure 6F). It is related to various metabolic and response processes, especially in the context of stress and stimulus responses like phosphorus and phosphate metabolism, response to external biotic stimulus, response to stress and biotic stimuli, apoptosis, oxylipin biosynthesis and lipid oxidation. The presence of stress response-related terms suggests that this module may play a role in adapting to environmental challenges.

#### Modules M26, M37, M81, M93, and M105

Present a correlation with RUBISCO, total chlorophyll, chlorophyll a, chlorophyll b, and PEPCASE (Figure 5). These modules are enriched in GO terms related to plant-type cell wall organization or biogenesis, negative regulation of growth, auxin polar transport, cellular component organization, cell wall polysaccharide metabolic process, hemicellulose metabolic process, peptide metabolic process, regulation of gene expression, and defense response. In contrast, modules correlated with PEPCASE (Figure 5) are associated with various combinations of metabolites and nitrogen conditions. No modules correlate with both genotype and nitrogen condition independently (Figure 5). However, module M123 shows significant correlation with the interaction between nitrogen condition and genotype. M123 is enriched in transcript groups involved in methylation and defense response. The absence of modules correlating nitrogen condition and genotype and the interaction between both, suggests genetic mechanisms underlying nitrogen use efficiency (NUE) differ from those governing the plant’s response to nitrogen availability. Alternatively, the results may reflect more complex interactions, such as non-monotonic relationships, that correlation analysis does not capture.

### Transcription associated proteins (TAPs) and nitrogen response

We found 39 TAPs belonging from families MYB/MYB-related: 57.4%, bHLH: 10.6%, CCAAT: 8.5%, WRKY: 6.38%, ARF and TRAF: 4.25%, Others: 2.12 %. The MYB/MYB-related is overrepresented (p value < 0.01). MYB family members were associated to Nitrogen-responsive sorghum genotypes (Gelli et al. 2014) and among DEGs of nitrogen tolerant sugarcane genotypes (Yang et al. 2019) These TAPs had different expression patterns, with some responding to genotype or nitrogen condition and some to both (Figure 7A).

**Figure 7.**
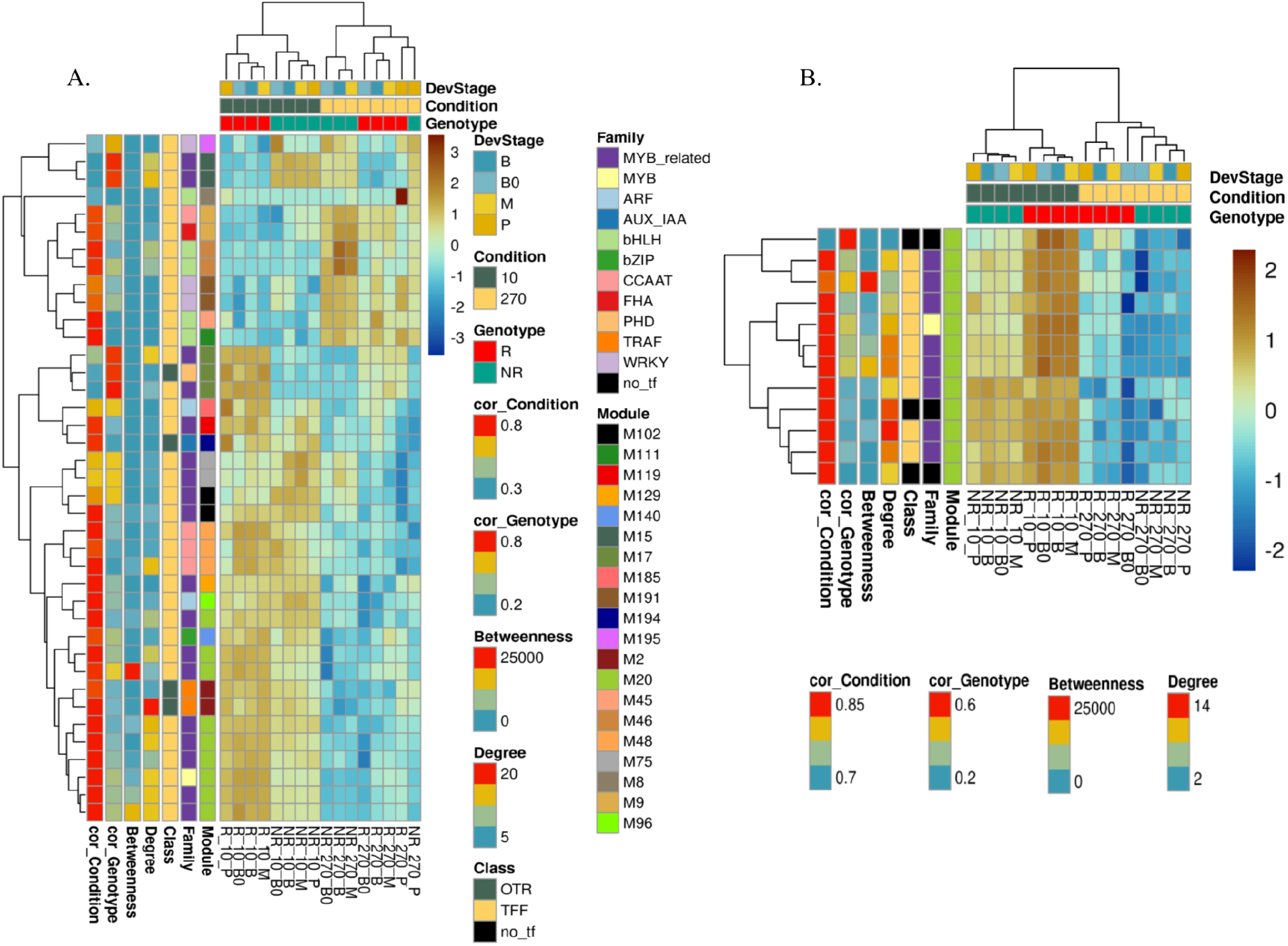
TAPs and module 20. A) TAPs expression pattern in the network. B) Expression pattern of module 20

Notably in module 20, 75% of transcript groups present are TFs from MYB or MYB-related families and they slightly respond to both genotype and condition. These factors were activated in low nitrogen conditions, more in the R genotype than in the NR genotype. Several transcript groups in this module show high (in the top 10% of highest in all network) degree and betweenness (Figure 7B) including SCA3__SP803280_c111248_g1_i2 with high betweenness, its annotated with transcription *cis*-regulatory region binding, cellular response to potassium, ion regulation of potassium ion transport, response to chitin, and the SCA3__SP803280_c108399_g1_i1 gene with high degree and annotated with protein ubiquitination, response to salt stress. The high degree and central betweenness of these transcripts indicate high centrality of this module in the network and possibly its biological importance orchestrating a core response to low nitrogen availability in both genotypes. Module 20 appears to be focused on processes related to development, metabolism, and regulation of biological functions, with a particular emphasis on floral development, reproductive processes, and regulatory mechanisms governing organismal development and senescence.

## Conclusion

This study provides novel insights into the molecular mechanisms underlying nitrogen utilization efficiency in sugarcane through gene co-expression network analysis. Our results revealed distinct transcriptional responses between responsive (R) and non-responsive (NR) genotypes, with 2,723 differentially expressed transcript groups identified across various leaf segments and nitrogen conditions. The responsive genotype showed enhanced expression of transcript groups involved in carbon metabolism and defense responses under high nitrogen conditions, while the non-responsive genotype prioritized photosynthesis and iron stress response pathways. Network analysis identified 199 distinct modules, with 44 modules correlating to nitrogen condition and 20 to genotype, suggesting complex and independent genetic mechanisms governing nitrogen response and genotype-specific traits. Notably, Module 20, enriched in MYB/MYB-related transcription factors, emerged as a potential key regulator of nitrogen response in both genotypes. The identification of novel candidate transcript groups, such as the uncharacterized NewTr2475430.gen in Module 15 and N_R__TRINITY_DN7789_c0_g2_i2.gen in Module 17, provides promising targets for future functional studies aimed at improving nitrogen use efficiency in sugarcane. These findings advance our understanding of nitrogen metabolism in sugarcane and offer valuable insights for breeding programs focused on developing more nitrogen-efficient varieties.

## Acknowledgements

This work was partially funded by the National Council for Scientific and Technological Development (CNPq), through a Research Productivity Fellowship awarded to DMRP (Grant #310080/2018-5) and a Universal Research Project granted to LM (Grant #442529/2014-7). Additional support was provided by the São Paulo Research Foundation (FAPESP) (Grants #2012/23345-0 to LM and #2013/15576-5 to MM). This study was also financed in part by the Coordenação de Aperfeiçoamento de Pessoal de Nível Superior – Brasil (CAPES) – Finance Code 001, through a PhD fellowship awarded to JMMP.

## Notes

https://figshare.com/projects/NitrogeGene_co-expression_networks_highlight_key_nodes_associated_with_ammonium_nitrate_response_in_sugarcane/242225

### Competing Interest Statement

The authors have declared no competing interest.

